# Rapid integration of artificial sensation

**DOI:** 10.1101/2025.04.14.648830

**Authors:** Samuel Senneka, Maria C. Dadarlat

## Abstract

1

Humans rely on both proprioceptive and visual feedback during reaching, integrating these two sensory streams to improve movement accuracy and precision [1, 2]. Patients using a Brain-Machine interface (BMI) will similarly require artificial proprioceptive feedback in addition to vision to finely control a prosthesis [3, 4]. Intracortical microstimulation (ICMS) elicits sensory perceptions that could replace the lost proprioceptive signal. However, some learning may be required for encoding artificial sensation [5], as current technology does not give access to neurons with all of the desired encoding properties [6]. We developed a freely-moving mouse behavioral task in which to test learning and integration of artificial sensory information. Five mice were implanted with a 16-channel microwire array in primary somatosensory cortex. Mice were trained to navigate to randomly-selected targets upon the floor of a custom behavioral training cage. Target location was encoded with visual and/or patterned multi-channel ICMS feedback. Mice received multi-modal feedback from the beginning of training of the behavioral task, achieving 75% on multimodal trials after approximately 1000 training trials. Mice also quickly learned to use the ICMS signal to locate invisible targets, achieving 75% proficiency on ICMS-only trials when tested. Critically, we found that performance on multimodal trials significantly exceeded unimodal performance (vision or ICMS), demonstrating that animals rapidly learned to integrate natural vision with artificial sensation.

**Significance:** Multisensory integration of visual and proprioceptive information facilitates accurate and precise movements. Intracortical microstimulation (ICMS) elicits perceptions that could supplement visual information for patients controlling a prostheses. Here, we developed a freely-moving mouse behavioral task to examine how ICMS can be used to encode multi-variable task-relevant information. Mice implanted with a cortical microwire array were trained to interpret patterned multi-channel ICMS to navigate to targets upon the floor of a custom behavioral training cage. Mice quickly learned to use the ICMS signal to locate invisible targets and integrated the artificial signal with natural vision, improving task performance. This protocol can be applied to efficiently develop and test algorithms to encode artificial proprioception for neural prostheses.

## 3 Introduction

The native human somatosensory system has access to an array of complex, dynamic inputs whose informational content is constructed from the activation of sensors distributed throughout the body [7]. The neural circuits underlying somatosensory processing remain plastic into adulthood, rapidly adapting to the statistics of sensory inputs: expanding the somatotopic representation of task-relevant information from the fingertips [8], developing novel multisensory responses to paired multimodal stimuli [9], and shifting receptive fields to incorporate extracorporeal objects such as tools [10]. Similarly, cortical neurons plastically adapt their response properties when directly activated by electrical currents (intracortical microstimulation; ICMS): expanding local somatotopic representations in response to peristant stimulation [11], altering functional connectivity between neurons [12], and shifting motor neuron projection fields [13]. Humans and animals can perceive neural activity elicited by artificial neuromodulation[14–20], making ICMS a promising tool for encoding artificial sensation to replace lost sensory function [21, 22] and to provide closed-loop feedback for neural prostheses such as Brain-Machine Interfaces (BMIs; e.g., [23]).

Animals learn to detect and interpret distinct spatial and temporal patterns of electrical stimulation across an array of electrodes, ultimately using multi-channel ICMS to guide perception and movement [17, 24–32]. This is particularly important for BMIs, which establish an artificial sensorimotor system [33–37]. Just like in natural sensorimotor function [1], BMI patients will require artificial somatosensation for fine control of a prosthesis [3, 4], which can be integrated with visual information to form a precise multisensory estimate [2, 38]. Modern neural interfaces enable high-dimensional neuromodulation via multi-channel ICMS — passing small electrical currents through chronically-implanted electrodes to mediate an “artificial sensation” [14, 15, 17, 39–45]. The prevailing philosophy is to pattern ICMS to evoke neural activity that closely resembles activity evoked by natural sensation. However, stimulation-evoked activity is state-dependent, and the required algorithms are computationally intensive, difficult to scale, and presume a known neural code [46– 50]. Additionally, there may be limited access to the population of neurons encoding the desired sensory variables [6, 41].

An alternative is to re-conceptualize the engineering challenge for artificial proprioception by replicating the information encoded by natural proprioception rather than replicating a specific pattern of neural activity. For example, proprioceptive information in primary somatosensory cortex arguably provides feedback on distance and direction of a movement [7, 51, 52]; Figure 1A). These two variables can be adopted to describe a different type of movements: navigation to a target in an open field (Figure 1B). Even if proprioception is instead encoded as a set of joint angles in primary somatosensory cortex [53], the goal should be to encode that set of variables (joint angles) rather than replicate specific neural activity patterns.

**Figure 1.**
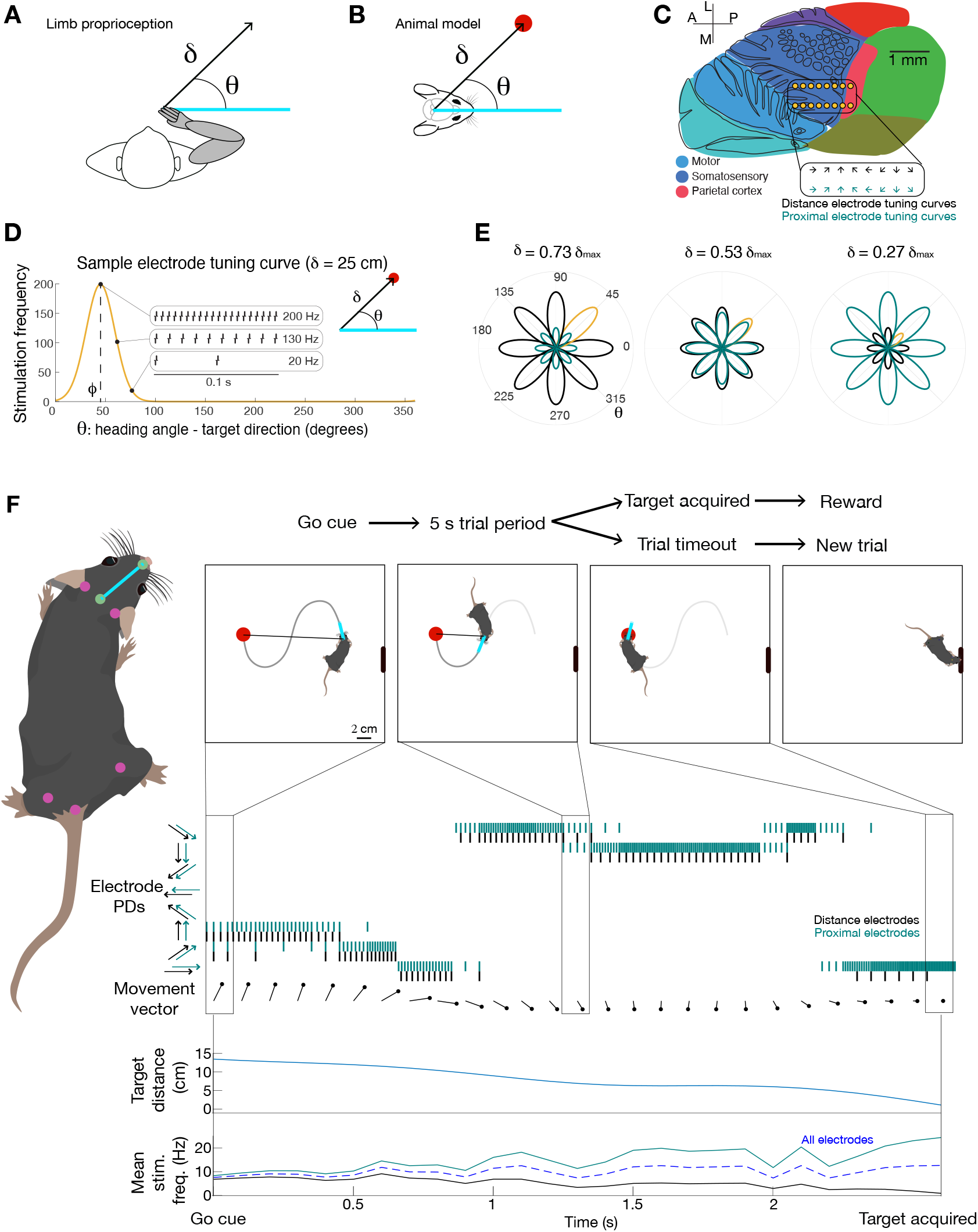
Encoding multi-variable artificial sensation via spatiotemporal patterned electrical microstimulation. **A.** Artificial sensation from a prosthetic limb encodes the distance (*δ*) and direction (*θ*) of movements. **B.** Animal model for artificial sensation: ICMS encodes the distance (*δ*) and direction (*θ*) between the animal’s heading and a rewarded target location. **C.** A sixteen-channel microwire array is implanted unilaterally over mouse primary somatosensory cortex (S), anatomically close to parietal (P), motor (M), and visual (V) cortices. **D.** Sample single electrode tuning curve with preferred direction *φ* shows stimulation frequency as a function of *θ* for fixed *δ*. **E.** Tuning curves for all sixteen electrodes as a function of target distance (*δ*) for distance (black) and proximal (green) electrodes. The yellow tuning curve shows the sample electrode in (D). **F.** A simulated behavioral trial, showing animal heading and movement trajectory (top). Left: animal pose is tracked in real-time within the cage (pink and green dots). Pose is used to calculate heading direction (vector between the nose and midpoint between the ears). Heading direction is used to calculate *δ* and *θ*. Stimulation parameters are updated at 10 Hz. Right: simulated stimulation patterns across “proximal” and “distance” group electrodes as a function of the movement vector across time (s). At bottom: mean stimulation frequency across the groups of proximal, distance, and all electrodes as a function of encoded movement vector during the trial.

Freely-moving rodents learn to use ICMS to navigate through an open field or to a reward port [26, 27, 54–57]. Nevertheless, movements guided by artificial sensation have so far failed to be as direct and precise as those guided by natural sensory signals [58]. The major questions from the perspective of encoding artificial sensation then are: 1) how to encode sensory information precisely, 2) how to encode multiple variables, and 3) what are the requirements for artificial sensation to be integrated with natural sensory information to improve artificial sensorimotor function? To address these questions, we developed a rapid and robust protocol by which to study encoding and integration of artificial sensation in freely moving mice. We showed that artificial sensation can be integrated with natural vision, and that integration is learned rapidly and robustly across animals. These results demonstrate the surprising flexibility of neural circuits in adapting to and integrating novel (artificial) forms of sensory information, which is an important consideration in the development of brain-computer interfaces.

## 4 Results

Five C57BL/6J mice were trained to navigate to a 2-cm diameter round target located on the floor of a custom behavioral training cage. The target could be set to one of nine possible locations, forming a three by three grid upon the cage floor (Suppl. Figure 1). The target location was made known to the mice via two sensory modalities: 1) vision (a circle upon the floor), and/or 2) multi-channel ICMS. The ICMS signal encoded a two-dimensional vector specifying 1) the angle between the mouse’s heading and the target (*θ*) and 2) the distance between the mouse’s head and the center of the target (*δ*; Figure 1B). The ICMS signal encoding of target location imitates the cosine tuning of neurons in sensorimotor cortices to the direction of limb movements [52, 59, 60]. To reduce the saliency of the visual target, it was dim (6 lux) and red (630 nm). Although mice are dichromats (they have no red cones in the retina), they are not blind to red light [61]. During each trial, the mouse’s head and body (Figure 1F) were tracked using DeepLabCut Live on a 60 Hz video feed from a camera mounted on the top of the behavioral training cage [62]. A behavioral trial was considered successful if the mouse’s head entered the target during a 5-10 s trial period.

### Encoding multi-variate artificial sensation

The artificial sensory signal was encoded via spatiotemporally patterned ICMS (Figure 1D-F). The basic unit of electrical stimulation is a biphasic pulse of current (Figure 1D), and the effect of stimulation on nearby neurons depends on the parameters used for stimulation, including the current amplitude, location of stimulation, and the frequency at which pulses are delivered [15, 40, 49, 63–65]. To encode the location of the target relative to the mouse’s head, each electrode was assigned a tuning curve in the form of a von Mises distribution centered on a “preferred direction” (*φ*) (Figure 1D), equally spaced around a circle from 0 to 2*π* (Figure 1E). The direction of the target relative to the mouse’s heading could be decoded from the relative stimulation frequencies across electrodes (Figure 1D,F). The distance between the mouse’s head and the center of the target (*δ*) scaled the amplitude of the tuning curve (maximum stimulation frequency; Figure 1E). To maintain a robust signal at all distances, electrodes were split into “proximal” and “distance” groups.

The proximal group tuning curves were scaled by (1 − *δ/δ_max_*)), increasing average stimulation frequency as the mouse approached the target. The distance group were scaled by (*δ/δ_max_*), decreasing stimulation frequency as the mouse approached the target. The distance between the target and the mouse’s head could therefore be decoded from the ratio of stimulation frequencies between the proximal and distance electrodes (Figure 1E,F).

### Learning artificial sensory information via multimodal stimulation

Animals received multimodal sensory information (both dim visual and ICMS encoding of target location) from the first day of training on the behavioral task. Mice quickly learned the multimodal behavioral task (Figure 2A), locating the target on at least 75% of trials within the first seven days of training. A level of 75% is often used as threshold for proficiency on a behavioral task [66]. Further training did not improve performance, as task completion on the last three days of testing was not statistically different from the first three days following proficiency (p ≥ 0.05, Wilcoxon rank-sum test). In both these early and late sessions, task completion was significantly better than the threshold set for proficiency (75%; p ≤ 0.05, t-test). Therefore, animals quickly learn and stabilize performance on this behavioral task.

**Figure 2.**
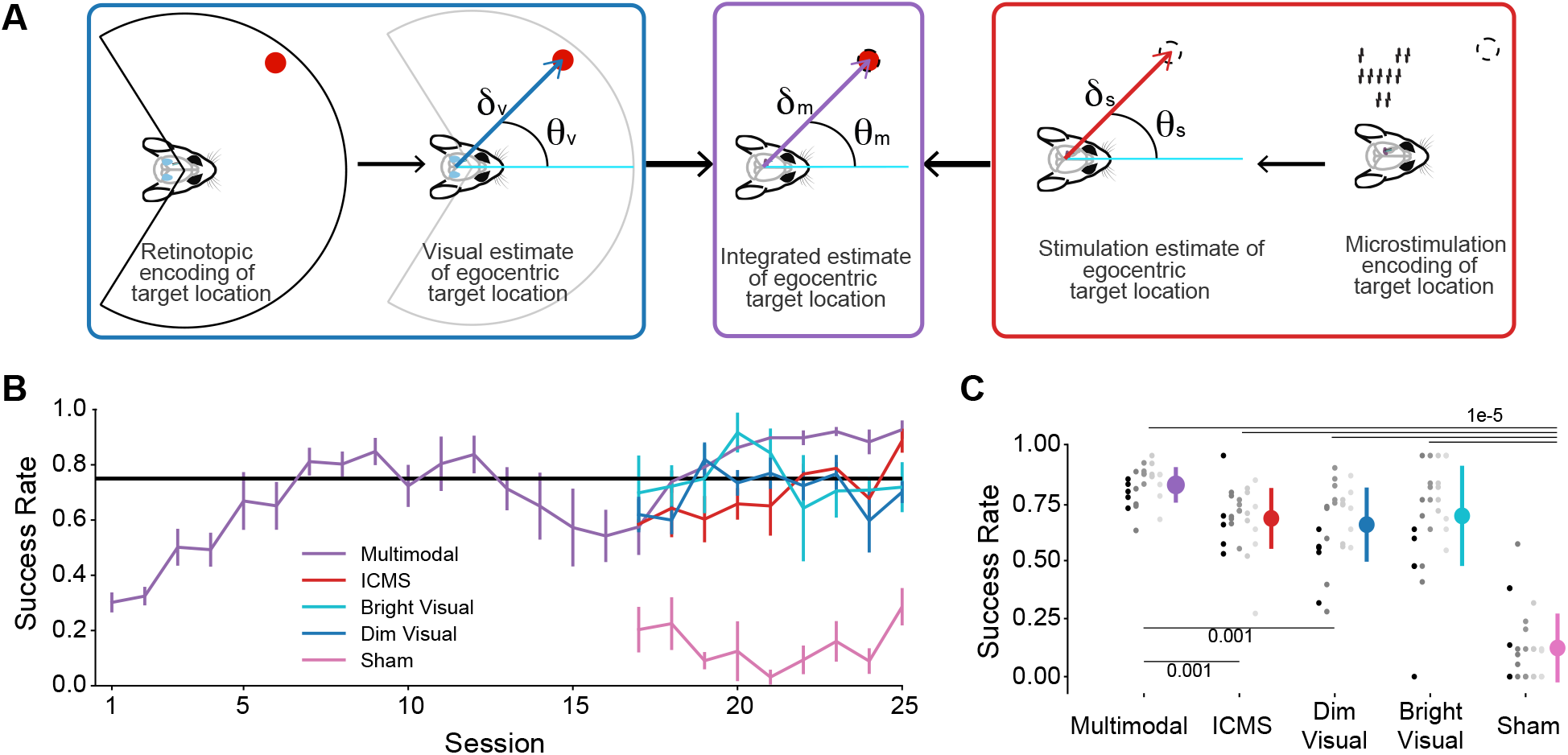
Learning to integrate artificial sensation. **A.** Computational requirements for multisensory integration of natural vision with artificial ICMS-based sensory encoding (*δ_m_,θ_m_*) includes combining visual estimates of distance and direction (*δ_v_,θ_v_*) with those estimated from the ICMS signal (*δ_s_,θ_s_*). **B.** Average and standard deviation of single-day success rates on each trial type across five animals. The noticeable dip around days 14-18 reflects poor performance for three of the mice after a break in training (day 13). **C.** Average success rates across trial types for all mice on the last five days of training. Each gray circle shows the performance of a single mouse on a single session. Error bars show std computed across all mice/sessions. Bars on the plot denote significant differences, with the p-value showing the outcome of a Wilcoxon sign-rank test, using a Holm-Bonferroni correction for multiple comparisons.

Once the mice were proficient on multimodal trials, we introduced a series of probe trials intended to test learning and integration of the ICMS signal. To test performance with the ICMS signal, ICMS probe trials were presented on 7.5 − 15% of trials per session, during which the target location was invisible to the mouse. We also included vision-only probe trials that could have dim or bright visual target trials in order to directly compare behavioral performance guided by natural and artificial sensation (7.5 − 15% and 2.5 − 5% of trials). Probe trials also included a sham condition, where no sensory information was provided about target location and therefore showed success rates achievable by chance (2.5 - 5%).

To quantify behavioral performance beyond success rate, we also calculated additional single-trial metrics: time-to-target, per-trial path efficiency, movement speed, and angular dispersion. The metrics were calculated for the last five behavioral sessions, at which point performance on multimodal trials had plateaued. These five stable sessions were used to strengthen the statistical comparisons of behavior guided by different sensory conditions. To ensure a fair comparison between vision and ICMS trial types, all analyses were performed on cropped trajectories — from the point the animals first faced the target and were less than 25 cm away from the target center. This was done to ensure that the trial statistics were not being calculated for periods during which ICMS or multimodal trials would provide additional information not present during vision trials, i.e., while the animal could not see the target and therefore could not make directed movements towards it.

Mice clearly learned to use the ICMS signal to locate the invisible targets within the training cage. Animals had stable average performance on ICMS trials, fluctuating around the 75% level set for proficiency (Figure 2B,C). In fact, behavioral performance guided by ICMS was qualitatively (Figure 3A) and quantitatively (Figure 3B-E) similar to behavioral performance when navigating to visual targets (p ≥ 0.05, Wilcoxon rank sum test). Under all sensory conditions, mice completed significantly more trials than chance (Sham), suggesting that mice are making directed movements towards the target on all trial types (p ≤ 1*e* − 5; Wilcoxon rank sum test, Holm-Bonferroni correction for multiple comparisons). Based on these results, we conclude that animals quickly learn to decode task-relevant sensory information from the ICMS signal and that decoding is stable across time.

**Figure 3.**
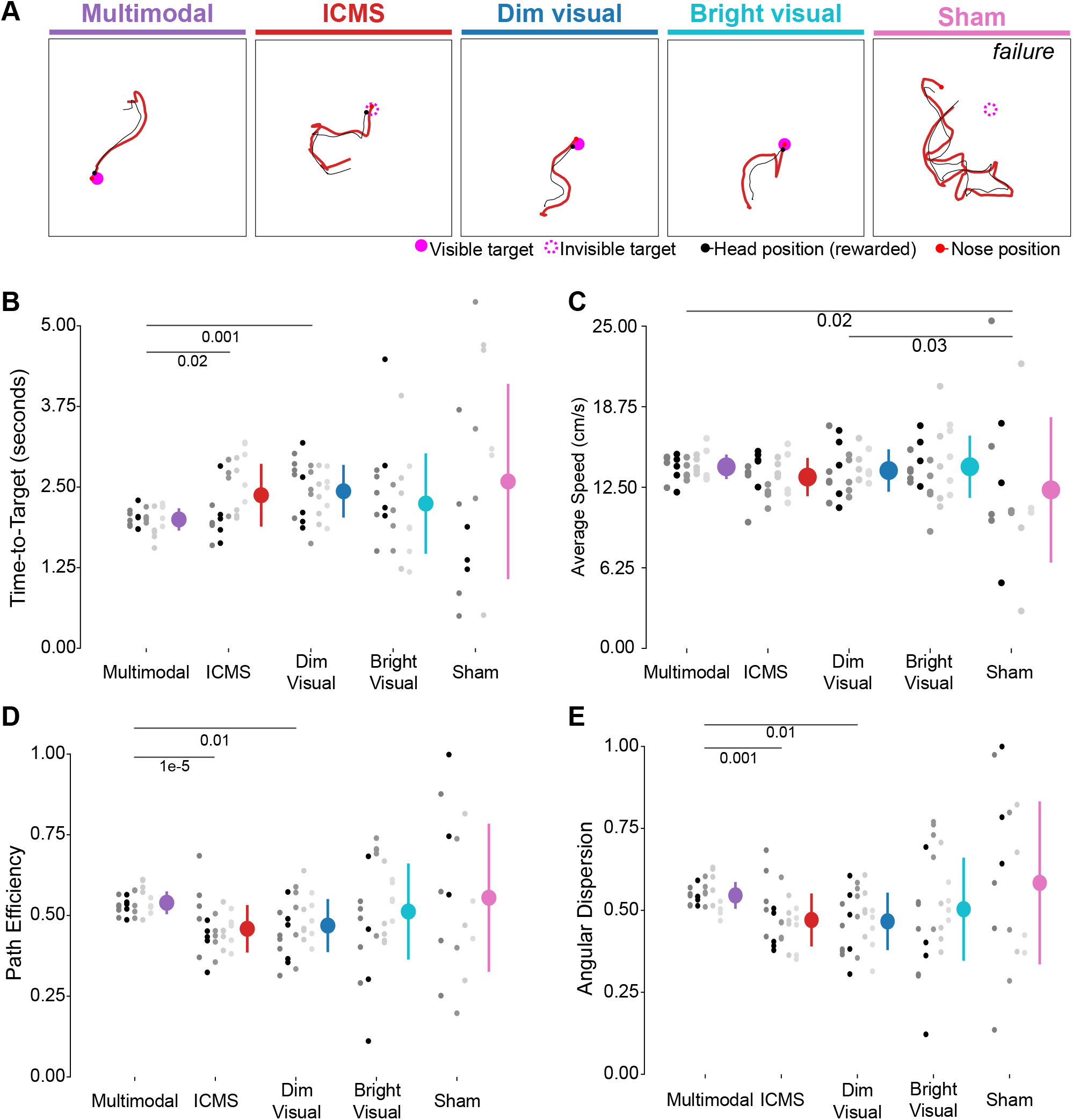
Comparison of task trajectories guided by natural, artificial, and multimodal sensation (p-values obtained with Wilcoxon rank-sum with Holm-Bonferroni corrections for multiple comparisons). Error bars in panels (B)-(E) show mean and STD across mice. Colors distinguish trial types, gray-scale dots represent individual mouse performance, and are consistent across all panels. All metrics are computed only for successful trials in the last 5 sessions of training. **A.** Sample single-trial trajectories for each trial type. **B.** Time taken to reach target (s). At the population level, mice were significantly faster performing the task with multimodal feedback than either unimodal ICMS or visual feedback. **C.** Average speed of movement (cm/s). Mice were significantly faster on multimodal and bright visual trials than sham trials on average. While mice were on average slower on ICMS trials compared to multimodal trials, this difference was not significant. **D.** Path efficiency. At a population level, mice took straighter paths on multimodal trials than ICMS or dim visual trials. **E.** Angular Dispersion. As with path efficiency, this metric provides a measure for how direct of a path the mice took to the target on successful trials. Statistical analyses indicate that mice took significantly straighter paths on multimodal trials than ICMS or dim visual trials at the population level.

### Integration of natural and artificial sensation

We hypothesized that, if natural visual information were being integrated with artificial sensory information (Figure 2A), behavioral performance on multimodal trials would be significantly better than on unimodal trials. This is in fact the case (Figures 2, 3A-E). Mice completed significantly more trials (Figure 2C) with more directed, efficient movements in multimodal than unimodal trials (Figure 3). Notably, mice were able to reach the target in a shorter period of time in multimodal than unimodal trials (Figure 3B), and moved faster in multimodal trials than in ICMS (but not visual) trials (Figure 3C). While ICMS immediately modulates neural activity [39, 50, 67], detection of ICMS may take longer than natural sensation [68, 69], slowing the animal’s movement. Another possibility is that the 10 Hz update rate of target location encoded via ICMS led to the animals moving more slowly to attend to the signal. In addition to movement speed, animals made more directed movements when guided by multimodal sensory information than by either ICMS or vision alone, as evidenced by shorter total path lengths and lower variability in movement direction in multimodal trials (Figure 3D,E). In conclusion, while there is prior evidence of multisensory integration of natural and artificial sensation [23, 30, 45, 58], our current work highlights the speed and flexibility by which integration can be learned.

### Decoding of direction and distance variables from ICMS

Theoretically mice could have high success rates with ICMS just by being more active in ICMS trials than in the sham trials. However, the statistical equivalence between ICMS and visual trials in path efficiency, angular dispersion, movement speed, and time-to-target suggest that target localization guided by ICMS is a deliberate process. Even so, we have not yet explicitly examined how well ICMS encodes each of *δ* and *θ* – the distance and direction between the mouse’s current heading and the target (Figure 1B). This question is challenging to address given the circuitous trajectory taken by the animals even to visual targets (Figure 3A). Therefore, we used behavioral performance on visual trials to estimate the intrinsic variability of mouse trajectories. Additional variability could then be assigned to an inability to extract task-relevant information about target location.

To estimate decoding of movement angle and distance from the ICMS signal, we first separated singletrial movement trajectories into discrete sub-movements based on changes in velocity [70]. The distinct sub-movements allows us to quantify changes in heading and distance relative to the target in response to sensory information available at the onset of the movement (Figure 4A). At the onset of movement, the mouse is at some location (*θ_onset_, δ_onset_*) relative to the target (Figure 4A, top left). If the animal can perfectly decode both direction and distance from the ICMS signal, an optimal movement would take the animal directly to the target, such that the change in angle over the sub-movement (the “estimated angle”), calculated as *θ*_onset_ − *θ*_offset_ = *θ*_onset_), could be compared to the true angle — the angle at movement onset. Similarly, the distance traveled in that sub-movement could be taken as the animal’s estimate of the true distance — the distance at movement onset. Only trials in which distinct sub-movements could be identified were included in this analysis. Furthermore, to remove noise from movements that were really just small twitches, only sub-movements that were of at least 200 ms duration were considered. Finally, this analysis was conduced on cropped trials, once the mice were facing the target, to make sure that the fist sub-movement was not just the mice rotating away from the reward port on the side of the behavioral training chamber.

**Figure 4.**
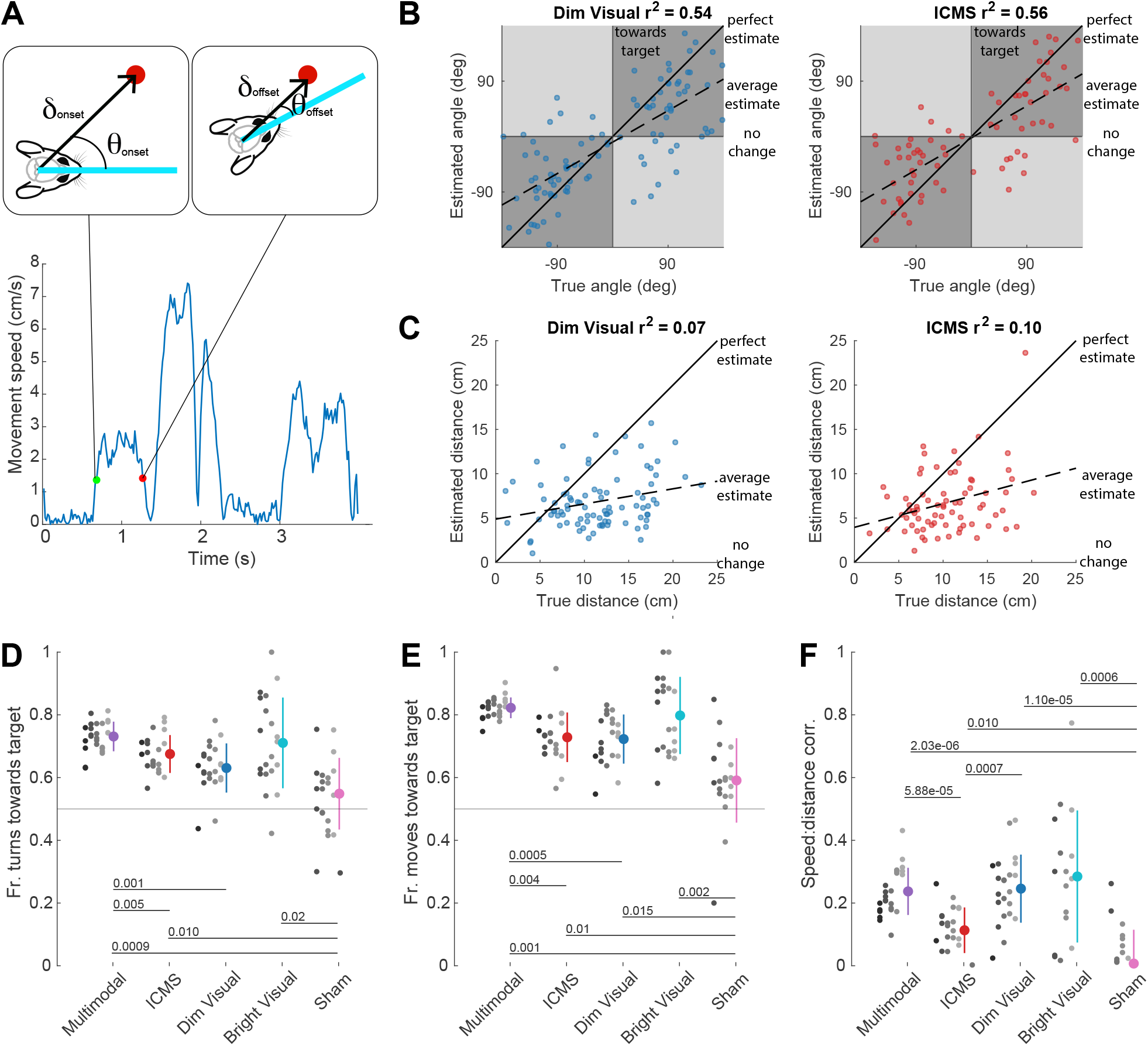
Estimation of relative target angle and distance from the ICMS signal. **A.** Example of identifiable sub-movements from the absolute radial speed of the mouse on a single trial. We use the first sub-movement of each trial (cropped to the point that the mouse first faces the target) to infer the animal’s estimate of the direction (*θ*) and distance (*δ*) between its head and the target at the beginning of the sub-movement. Estimated angle is calculated from the change in angle between submovement onset and offset (*θ*_onset_ and *θ*_offset_). Similarly, distance estimation is inferred from the change in distance between sub-movement onset and offset (*δ*_onset_ and *δ*_offset_). Data comes from all five mice. **B.** Decoding *θ* on each trial type. Each circle represents the first sub-movement of Dim Visual trials (left) and ICMS trials (right) in trials with clear changes in velocity, as shown in (A). The black diagonal line represents unity – a perfect estimate of angle. No rotation would yield an estimated angle of 0*^o^*. A linear regression was fit between the true angle (x-axis) and the estimated angle (y-axis) on this dataset (dashed line). The *r*^2^ value of the fit is shown in the title. **C.** Data as in (B), but examines estimates of target distance: *δ*. The black diagonal line represents unity – a perfect estimate of distance. No movement would yield an estimated angle of 0*^o^*. A linear regression was fit between the true target distance (x-axis) and the estimated distance (y-axis) on this dataset (dashed line). The *r*^2^ value of the fit is shown in the title. **D.** Fraction of turns towards the target, calculated by splitting single trials into 6 cm segments (the median of sub-movement distance calculated in (C)). These values are calculated only for correctly completed trials, including in the shamcondition. P-values show statistically significant differences in facing between trial types using a Wilcoxon rank-sum test with a Bonferroni correction for multiple comparisons. **E.** Fraction of movements towards the target, calculated by splitting single trials into 6 cm segments (the median of sub-movement distance calculated in (C)). These values are calculated only for correctly completed trials, including in the sham condition. P-values show statistically significant differences in facing between trial types using a Wilcoxon rank-sum test with a Bonferroni correction for multiple comparisons. **F.** Single-trial correlations between movement speed and distance to the target is used to indirectly infer the mouse’s ability to calculate relative distance, *δ*, across trial types. These values are calculated only for correctly completed trials, including in the Sham condition. P-values show statistically significant differences between trial types, calculated using a Wilcoxon rank-sum test with a Bonferroni correction for multiple comparisons.

For both visual and ICMS trials, we found that the estimated angle was linearly related to the true angle, although mice tended to underestimate the true angle (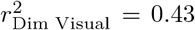 and 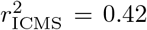; Figure 4B). Even so, the strong correlation between the two suggests that mice were able to extract target direction information from ICMS similarly well to vision. In contrast, estimations of target distance were generally poor, with little relationship to the true target distance at sub-movement onset (Figure 4C). Perhaps this poor correlation is not surprising, because the use of distinct sub-movements to reach the target suggests that the animal has low certainty about the target location on those trials and is stopping to accumulate more information.

To further quantify how well *δ* and *θ* can be decoded from the ICMS signal, we performed three additional analyses. First, we hypothesized that, on average, the mouse’s movement should be turning its head towards the target. To test this idea, we separated each trial into segments of 6 cm (the median of the distribution of distances traveled during distinct sub-movements that lasted longer than 200 ms). We then calculated the fraction of these 6 cm segments during which the mouse was turning towards rather than away from the target direction. We found that, just as the behavioral metrics described in prior analyses, the mouse made just as many turns towards the target during ICMS trials as to visual trials (Figure 4D). Furthermore, mice made more turns towards the target in multimodal trials than unimodal ICMS or Dim Visual trials (*p* ≤ 0.005, Wilcoxon sign-rank test), once again supporting the notion that artificial and natural sensation are integrated.

Fraction of turns towards the target primarily examines decoding of directional information. To examine decoding of distance information, we modified the analysis to ask on what fraction of movement segments did the mouse come closer to the target. If the mouse were moving randomly, as in the Sham condition, movements would only bring the animal closer to the target approximately half of the time. As before, we considered movement segments of 6 cm. A movement was considered to be “towards” the target if the radial distance to the target at the endpoint of the segment was smaller than the radial distance at the start of the segment. As before, performance with ICMS was similar to that with the Dim Visual signal, and performance with the multimodal signal significantly exceeded unimodal trials (Figure 4E).

Finally, we noted that mice slowed down as they approached the bright visual target, reflected by a positive correlation between target distance and movement speed (*ρ* = 0.28; Figure 4E). This behavior may reflect the well-known trade-off between the speed and accuracy of movements (Fitts’s law), suggesting that movements become slower and more precise as the animal knowingly approaches the target. Similarly, target distance and movement speed were positively correlated for ICMS trials (with correlations significantly greater than zero across trials, p = 5*e* − 8, t-test). However, the correlation between target distance and movement speed was lower for ICMS trials than for either dim visual trials or bright visual trials (p = 5.9*e* − 5 and p = 7*e* − 4, Wilcoxon rank sum test, using Bonferroni correction for multiple comparisons), suggesting that distance information was not as well encoded by the ICMS signal as by the visual signal.

## 5 Discussion

Our major finding is that freely-moving animals quickly integrate multi-variate artificial sensation encoded by patterned ICMS with natural vision to improve behavioral performance on multimodal trials relative to unimodal trials. Furthermore, although we used a learning-based approach towards encoding the artificial sensory signal, the mice ultimately perform just as well on unimodal ICMS trials as on visual trials, as measured by task success rates, path efficiency, path variability, movement speed, and trial duration. A closer analysis of single-trial trajectories suggested that animals learned to decode direction, and to a lesser extent, distance, from an ICMS signal, with accuracies comparable to natural vision. The speed and reliability of multisensory integration across animals emphasizes the flexibility of the nervous system in processing and adapting to novel forms of sensory information, including integrating information that is not available via natural sensors. Our results suggest that integration is a natural consequence of learning in the context of sensory-guided behavior [71]. We propose that this protocol may be used to rapidly study the optimal methods for multi-variate encoding of artificial sensation.

### Multisensory integration must be learned through experience

Multisensory integration is characteristic of natural sensory processing, and has been identified across a range of species, brain areas, sensory modalities, and behavioral tasks [2, 38, 72–79]. The process of multisensory integration can and must be learned through experience of paired sensory information [71, 80–82]. This can happen quickly – neurons can develop multisensory responses in naive adult animals in as few as 3,600 cross-modal exposure trials (six hours), even under anesthesia [80]. Once established, mulstisensory responses are stable, even without subsequent multisensory exposure [83]. Multisensory integration has also been shown behaviorally in the context of artificial sensation, even when stimulation is not biomimetic [23, 58]. However, when learning of integration was explicitly studied, it took an order of magnitude longer (60,000 trials) for integration of natural and artificial sensation in head-fixed macaques. In contrast, freely-moving mice integrated visual information with artificial sensation encoded by electrical stimulation after just 20 behavioral training sessions (roughly 3,000 trials or 2 hours of multimodal sensory experience). This speed may come from the simplification of the behavioral task (navigating to a visual target) and the improved ethological relevance of the behavior to the animal. While additional work will be needed to establish the neural basis of integration of natural and artificial sensory information, the speed at which it occurs suggests ready integration into natural sensorimotor circuits.

### Artificial proprioceptive information is required for precise control of a robotic limb

Artificial proprioceptive feedback from a prosthesis should encode the position and movement of the limb through space [7]. A major engineering challenge is thus to encode as many sensory variables as possible (feedback from the shoulder, elbow, hand, fingers, etc.) with as few electrodes as possible (e.g., 100 in the Utah array [84]). Therefore, refining algorithms used to encode artificial sensation to improve its reliability and precision is a critical area of research. Previous work in freely-moving rodents has demonstrated speed of learning of ICMS in this context, suggesting that this model is ideal for early development of encoding strategies for artificial sensation [26, 27, 54–57].

### How much learning is necessary for artificial sensation?

An informal distinction in the field of artificial sensation is whether or not neuromodulation is “biomimetic.” This term conveys the idea that an optimal form of artificial sensation is one that takes no learning, by which electrical stimulation can directly replace natural sensation in the context of behavioral perception and discrimination (e.g., [16, 18]). Intuitively, the nervous system is optimized to interpret and generate specific activity patterns, and movement away from those patterns requires neural plasticity[85–87]. Consequently, understanding how electrical stimulation can be patterned to evoke neural activity patterns similar to those induced by natural sensation is of interest [44, 46, 47, 88]. These efforts have been particularly successful in the field of artificial touch, where neurons have readily-identifiable receptive fields and electrical stimulation induces sensation on clearly demarcated parts of the body [17, 89, 90].

Achieving similar results for artificial proprioception is challenging, due to the complexity of the proprioceptive neural code [52, 59, 60] (but see [6, 91]). In general, any finite number of electrodes is unlikely to be able to precisely access and modulate neurons with all of the desired encoding properties [6]. Therefore, encoding the full range of artificial proprioceptive signals from even a single limb may require some level of learning from the subject. Encouragingly, the sensory perceptions elicited by peripheral nerve microstimulation shift to align to prosthesis sensor locations [5]. However, persistent microstimulation may be required to engage plasticity, as transient stimulation shows stable evoked perceptions over long periods of time [92]. Our approach investigates an extreme circumstance: what if almost *everything has to be learned*? Addressing this question sets a lower bound for behavior guided by electrical stimulation and demonstrates the extent to which learning and plasticity can be engaged by neural prostheses. That said, as mentioned earlier, our design of stimulation patterns was inspired by observations regarding neural encoding of natural sensation. Specifically, we assigned electrodes gaussian tuning curves to movement direction, which imitate the tuned responses of single neurons to motion [52, 93, 94]. This may be an important consideration for encoding artificial proprioception.

While our analyses clearly suggest that mice can extract the encoded sensory variables from multi-channel ICMS, one confound is that ICMS could trigger a global state of arousal (e.g., [95]), making the animals move more, increasing the likelihood of finding the target by chance than in sham trials. However, given that there is no difference in path efficiency between ICMS and visual trials (Figure 3E), this scenario is unlikely. A contrasting concern is that ICMS may take longer to process than natural sensation, leading to slower movements than those guided by natural sensation. When a single electrode is used to encode a tactile sensation, the response time to report the stimulus is sometimes, but not always, longer for ICMS encoding than for vision or touch [68, 69]. However, response times speed up if stimulation is delivered with higher intensities or across more electrodes [96]. In this study, the tuning curves of the electrodes were narrow enough that typically only a pair of electrodes were stimulated at a time (Figure 1D-F). In our experiments, there was no difference in movement speed or time-to-target between visual trials and ICMS trials. Therefore, the ICMS signal seems to be equally informative as natural vision in this task. More work will be needed to address the question of whether it will be just as informative in the context of BMIs, but our results are encouraging.

## 6 Methods

### 6.1 Mice

All procedures were approved by the Institutional Animal Care and Use Committee at Purdue University. Five C57BL/6J mice (Jackson Labs #000664) aged 3-4 months were used in these experiments (4 males, 1 female). Mice were housed in groups of no more than five per cage in a temperature and humidity-controlled room on a 12-hour light/dark cycle with *ad libitum* access to food and water prior to surgery. Mice were single-housed following surgery for the duration of the experiments. During training, mice had free access to citric acid water (0.5-5%) within their home cages [97]. In the behavioral training cage, mice received a 0.1 mL strawberry Nesquik reward for each successful trial.

### 6.2 Surgical Procedures

Each mouse underwent a chronic microwire array implantation surgery prior to behavioral training. 16channel microwire arrays (TDT OMN1010-16) were implanted over the forelimb/hindlimb region of mouse primary somatosensory cortex. This region was identified using stereotaxic coordinates (Paxinos and Franklin’s the Mouse Brain in Stereotaxic Coordinates).

To prepare for surgery, each was induced to anesthesia using 1-3% isoflurane and maintained at 1-2% to areflexia. Ophthalmic ointment was applied to each eye. Mice were then given subcutaneous doses of meloxicam (10 mg/kg) and dexamethasone (100 mg/kg) prior to the start of surgery. Next, 0.2% lidocaine (.03 mL) was injected cutaneously on the scalp and the scalp was resected.

Prior to electrode array implantation, a ∼ 1 x 2.5 mm craniectomy was made in the right hemisphere over the posterior portion of primary somatosensory cortex and the anterior portion of posterior parietal cortex. Electrode arrays consist of 16-channel microwire arrays in an 8x2 layout. Each electrode is 2 mm long. Within each row of the array (8 electrodes), electrodes are spaced 250 *μ*m apart, and the two rows are 500 *μ*m apart. Electrode arrays were implanted to a depth of 500 *μ*m using a digital stereotax (Stoelting). Following array implantation, a thin layer of silicone elastomer (KwikSil) was used to cover the exposed brain, and the array was secured to the skull with dental cement (Metabond).

To implant ground and reference wires, two burr holes were created in the contralateral hemisphere. Using these burr holes, silver ground and reference wires were implanted just underneath the skull. On three of the animals, a two mm 0-80 screw was also placed in the temporal ridge contralateral to implantation site to secure the implant. Additional dental cement was applied to secure the array, ground and reference wires, and screw.

Mice were treated post-operatively with meloxicam (10 mg/kg) and dexamethasone (100 mg/kg) for three and seven days post-op, respectively, and were given five to seven days to recover before training began.

### 6.3 Behavioral training

#### Behavioral training chamber

The behavioral task required mice to locate a target that could appear in one of nine locations upon the floor of the behavioral training chamber. The chamber was a custom-designed and fabricated rectangular chamber (24 cm x 25.4 cm) made of acrylic with a clear floor. Visual targets were generated using a 32x32 LED RGB matrix with a pitch of 6 mm (Adafruit Product ID: 1484) placed just underneath the cage floor. Visual targets were generated by turning on groups of LEDs using red light (630 nm) with an intensity of 6 lux for the visual target, and were chosen randomly on each trial. A reward port was placed upon the center of one wall, approximately 2 cm off the ground. Two speakers (Fostex FT17H Horn Super Tweeter) were mounted on opposing walls to provide auditory cues, including a start tone (20 kHz, 0.5 s duration) and a reward tone (8 kHz, 1 s duration). Neural stimulation and recordings were performed using a Ripple Neuro Grapevine processor and Ripple Neuro pico+stim headstage. The mouse’s position within the cage was recorded at 60 Hz with a camera mounted on the ceiling of the training chamber (Imaging Source monochrome camera DMK 37BUX287 used with an Imaging Source TPL 0420 6MP Lens). We used a trained DeeplabCut pose estimation model and the DeepLabCut-Live GUI to track the mouse’s movement on the real-time camera feed [62]. A custom python script was used in conjunction with DeepLabCut-Live to randomly select target regions, provide reward, and update stimulation parameters during trials.

#### Habituation

Animals were placed on 0.5-5% citric acid water within their homecage [97] and were habituated to the behavioral training chamber for three days prior to the start of behavioral training. Habituation consisted of 5-15 minutes sessions. During the habituation sessions, a reward port (Sanworks Product ID: 1009) was activated for a portion of every minute, alternating between an activation period (55 s) and a deactivation period (5 s). Reward port activation was indicated by a white LED on the port being illuminated and a tone being played at the onset of activation (1 second 8 kHz pure tone). If the mouse nose-poked the reward port during the 55 second activation period, the solenoid on the port was opened for 10 ms, delivering a drop of strawberry Nesquik. Subsequent nose-pokes did not provide reward until next activation period. Therefore, mice could receive a reward once every 60 s during habituation.

#### Training

Subjects were initially trained to navigate to multimodal targets. To make the trials easy to complete, the dim visual targets were large (7-9 cm diameter) and the trial duration was long (10-12 s). As subjects completed approximately ≥ 80% of trials on each condition, target size and trial duration were gradually reduced to the final values of 2 cm diameter visual targets and 5 second trials. The training process took approximately four session, using consistent decreases in target diameter of 2 cm/session and 1-2 seconds/per session.

#### Testing

Once mice performed well (≥ 70% success rates) on multimodal trials, four additional trial types were introduced: ICMS-only, visual-only (dim and bright), and sham (no visual or ICMS). ICMS-only explicitly tests decoding of the artificial sensory signal, sham trials test success rates that are achievable by chance (setting a lower bound on performance), and the visual trials test baseline behavioral guided by unimodal natural feedback when the signal is highly reliable (bright visual) or less reliable (dim visual). The first two mice were tested using a trial probability distribution of 80% multimodal, 7.5% ICMS only, 7.5% dim visual only, 2.5% bright visual only, and 2.5% sham. To gain a more precise estimate of unimodal behavior, three subsequent mice were tested using a trial probability distribution of 60% multimodal, 15% ICMS only, 15% dim visual only, 5% bright visual only, and 5% sham.

### 6.4 Intracortical Microstimulation

ICMS pulses were cathode-leading biphasic symmetric pulses with a phase length of 200 *μ*s and interphase interval of 250 *μ*s with a constant amplitude of 10-20 *μ*A. The frequency and location of stimulation were used to encode the vector describing the relative position between the mouse’s forward heading and the target. For electrode *i*, stimulation frequency at time *t* was set to

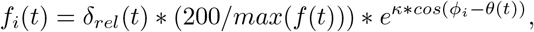

where *θ*(*t*) is the target direction relative to the animal’s heading, *φ_i_* is fixed for each electrode, *δ*(*t*) scales with distance to the target, and 200*/max*(*f* (*t*)) scales the curves to a maximum stimulation frequency of 200 Hz. *δ*(*t*) was modeled as *δ_rel_*(*t*) = *δ*(*t*)*/δ_max_* and *δ_rel_*(*t*) = 1 − *δ*(*t*)*/δ_max_* for distal and proximal encoding electrodes, respectively, with *δ_max_* = 26.5 cm (the diagonal distance across cage). Electrodes 1-8 (row 1) were set to be proximal encoding electrodes and electrodes 9-16 (row 2) were set to be distance encoding electrodes. Both electrodes located within each column of the 8 x 2 array had the same *φ_i_*, a value that acted as a “preferred direction” for the electrodes as it is the direction at which the electrode had the highest stimulation frequency for fixed *δ*.

### 6.5 Analysis

#### Statistics

Statistical analyses for unpaired samples were all performed using t-test or Wilcoxon rank sum test and corrected for multiple comparisons using the Bonferroni correction. Paired samples (such as comparing behavior of a single mouse on different trial types each day) were tested using the Wilcoxon sign-rank test.

#### Behavioral analysis

Trajectories taken during trials were cleaned prior to analysis due to noise in the DeepLabCut tracking algorithm. This cleaning was achieved by identifying time points with low probability of being correct (a measure provided by DeepLabCut) and averaging the value of the two surrounding, higher-probability points. For all metrics excluding success rate, only successful trials were included in statistical analyses.

##### Trial cropping

On a subset of trials, the mouse is not originally facing the target. To ensure a fair comparison between visual and ICMS trials, we cropped behavioral trajectories on each trial to the point at which the mouse is first facing the target, defined as the point at which the mouse heading (vector between the nose and back of the head; Figure 1) is within *± 123°* of the target direction, *δ*. Subsequent trial metrics (described below) were calculated using these cropped trajectories. We then quantified behavioral performance under each sensory condition with three behavioral measures: success rate, path efficiency, and time-to-target.

*Path efficiency* is calculated as the ratio between the direct distance to the target at beginning of the trial and the total path length traveled by the animal during the trial. The total path length is calculated as 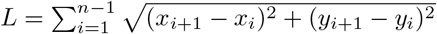 where *x* and *y* are the coordinates of the mouse’s location captured by DeepLabCut and *n* is the number of data points collected in a single trial. For example, a path efficiency of 1 indicates that the animal took a straight-line path to the target, whereas a path efficiency of 0.5 indicates that the animal traveled twice as far as the most direct path.

*Angular Dispersion* measures the spread of turning angles around an average direction of movement and provides a second measure of path tortuosity. This measure is calculated using the formula 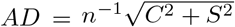 where 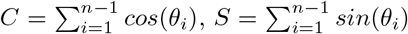, *θ_i_* is the angle of movement between successive positions along the trajectory of movement and n is the number of data points collected along the path. This formula results in a value from 0 to 1 with 0 being a completely distributed, circular path, and 1 being a straight path.

*Time-to-target* is measured as the total time between the start tone and the time the mouse’s head reaches the target. This provides a raw measure of how quickly mice find the target on average, regardless of starting distance from the target.

*Average Speed* is calculated as 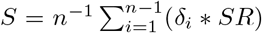 where *δ* is the distance between successive time points, SR is the sampling rate (60 Hz), and n is the number of data points collected in a single trial.

##### Estimation of angle and direction from sensory information

To quantify how well the animal could estimate the angle and direction of the target from each sensory modality, we isolated trials in which the velocity profiles had clearly identifiable bell-curves. A bell-curve shaped velocity profile was defined as one that had “onset” and “offset” points equal to baseline speed plus 0.5*σ_v_*, where *σ_v_* is the standard deviation of the cropped velocity trace. Velocity on each trial was calculated as the difference in position of the animal normalized by the inverse of the camera frame rate (1/60 Hz^−^1). Only movement segments of at least 200 ms duration were considered, to reduce noise from small, unintentional head movements. This analysis includes trials in which the mouse was moving when the go cue sounded, slowed down to identify the target location based on the available sensory information, and then resumed moving. On these trials, we used the first sub-movement following the first pause to assess decoding of *θ* and of *δ*.

##### Fraction of turns and movements towards target

We used two strategies to break up single-trial trajectories into sub-movements. As most trials did not have the distinct sub-movements evident in Figure 4A, trajectories were split by either distance or time.

To calculate the fraction of movements towards the target (to assess inference of *δ*), we split each trial into segments of 6 cm duration, which was the median of the movement distance calculated for the initial sub-movements. We then found the radial distance between the mouse’s head and the center of the target at the onset and offset of the movement segment. If the radial distance decreased from movement onset to offset, that segment was labeled as a move towards the target. The total number of movements towards the target was normalized by the total number of segments on that trial to calculate a single-trial fraction of movements towards the target. The same set of segments were used to infer decoding of *θ*. We calculated the relative angle to the target at the start and endpoint of each of these segments. That segment was considered a turn towards the target if the absolute value of the relative offset angle was smaller than the absolute value of the relative onset angle. The total number of turns towards the target was normalized by the total number of segments on that trial to calculate a single-trial fraction of turns towards the target.

## 7 Acknowledgments

The work described here was supported by NSF HDR grant 2117997, NIH T-32 grant 016853-06 for Purdue University’s Interdisciplinary Training Program in Auditory Neuroscience, and the Indiana CTSI Traumatic Spinal Cord & Brain Injury Research Program.

## References

[1] Philip N. Sabes. “Sensory integration for reaching. Models of optimality in the context of behavior and the underlying neural circuits”. In: Progress in Brain Research 191 (2011), pp. 195–209.

[2] Marc O Ernst and Martin S Banks. “Humans integrate visual and haptic information in a statistically optimal fashion.” In: Nature 415.6870 (2002), pp. 429–433. eprint: NIHMS150003.

[3] Sharlene N. Flesher et al. “A brain-computer interface that evokes tactile sensations improves robotic arm control”. In: Science 372.May (2021), pp. 831–836.

[4] A. J. Suminski, D. C. Tkach, A. H. Fagg, and N. G. Hatsopoulos. “Incorporating Feedback from Multiple Sensory Modalities Enhances Brain-Machine Interface Control”. In: Journal of Neuroscience 30.50 (2010), pp. 16777–16787.

[5] Ivana Cuberovic, Anisha Gill, Linda J. Resnik, Dustin J. Tyler, and Emily L. Graczyk. “Learning of Artificial Sensation Through Long-Term Home Use of a Sensory-Enabled Prosthesis”. In: Frontiers in Neuroscience 13.August (2019), pp. 1–24.

[6] Tucker Tomlinson and Lee E Miller. “Toward a Proprioceptive Neural Interface That Mimics Natural Cortical Activity”. In: Adv Exp Med Biol. 957 (2016), pp. 367–388.

[7] Benoit P. Delhaye, Katie H. Long, and Sliman J. Bensmaia. “Neural basis of touch and proprioception”. In: Compr Physiol. 8.4 (2018), pp. 1575–1602.

[8] G. H. Recanzone, M. M. Merzenich, W. M. Jenkins, K. A. Grajski, and H. R. Dinse. “Topographic reorganization of the hand representation in cortical area 3b of owl monkeys trained in a frequency-discrimination task”. In: Journal of Neurophysiology 67.5 (1992), pp. 1031–1056.

[9] Solaiman Shokur et al. “Expanding the primate body schema in sensorimotor cortex by virtual touches of an avatar”. In: Proceedings of the National Academy of Sciences of the United States of America 110.37 (2013), pp. 15121–15126.

[10] Angelo Maravita and Atsushi Iriki. “Tools for the body (schema)”. In: Trends in Cognitive Sciences 8.2 (2004), pp. 79–86.

[11] G H Recanzone, M M Merzenich, and H R Dinse. “Expansion of the cortical representation of a specific skin field in primary somatosensory cortex by intracortical microstimulation.” In: Cerebral cortex (New York, N.Y. : 1991) 2.3 (1992), pp. 181–96.

[12] James M. Rebesco, Ian H. Stevenson, Konrad P. Körding, Sara A. Solla, and Lee E. Miller. “Rewiring neural interactions by micro-stimulation”. In: Frontiers in Systems Neuroscience 4.August (2010), pp. 1–15.

[13] Andrew Jackson, Jaideep Mavoori, and Eberhard E Fetz. “Long-term motor cortex plasticity induced by an electronic neural implant.” In: Nature 444.7115 (2006), pp. 56–60.

[14] S. N. Flesher et al. “Intracortical microstimulation of human somatosensory cortex”. In: Science Translational Medicine 8.361 (2016), 361ra141–361ra141.

[15] Christopher L. Hughes et al. “Perception of microstimulation frequency in human somatosensory cortex”. In: eLife 10 (2021), pp. 1–19.

[16] Kevin J. Otto, Patrick J. Rousche, and Daryl R. Kipke. “Microstimulation in auditory cortex provides a substrate for detailed behaviors”. In: Hearing Research 210.1-2 (2005), pp. 112–117.

[17] Giacomo Valle et al. “Tactile Edges and Motion Via Patterned Microstimulation of the Human Cortex”. In: Science 387 (2025), pp. 315–322.

[18] Ranulfo Romo, Adrián Hernández, Antonio Zainos, and Emilio Salinas. “Somatosensory discrimination based on cortical microstimulation”. In: Nature 292.March (1998), pp. 387–390.

[19] Aamir Abbasi, Dorian Goueytes, Daniel E. Shulz, Valérie Ego-Stengel, and Luc Estebanez. “A fast intracortical brain-machine interface with patterned optogenetic feedback”. In: Journal of Neural Engineering 15.4 (2018).

[20] Mario Prsa, Gregorio L. Galiñanes, and Daniel Huber. “Rapid Integration of Artificial Sensory Feedback during Operant Conditioning of Motor Cortex Neurons”. In: Neuron 93.4 (2017), 929–939.e6.

[21] Léo Pio-Lopez, Romanos Poulkouras, and Damien Depannemaecker. “Visual cortical prosthesis: an electrical perspective”. In: Journal of Medical Engineering and Technology 45.5 (2021), pp. 394–407.

[22] Xing Chen, Feng Wang, Eduardo Fernandez, and Pieter R. Roelfsema. “Shape perception via a high-channel-count neuroprosthesis in monkey visual cortex”. In: Science 370.6521 (2020), pp. 1191–1196.

[23] G. Risso et al. “Optimal integration of intraneural somatosensory feedback with visual information: a single-case study”. In: Scientific Reports 9.1 (2019), pp. 1–10.

[24] Roy Lycke et al. “Low-threshold, high-resolution, chronically stable intracortical microstimulation by ultraflexible electrodes”. In: Cell Reports 42.6 (2023), p. 112554.

[25] Amy M. Ni and J. H R Maunsell. “Microstimulation Reveals Limits in Detecting Different Signals from a Local Cortical Region”. In: Current Biology 20.9 (2010), pp. 824–828. eprint: NIHMS150003.

[26] Eric E. Thomson, Rafael Carra, and Miguel A.L. Nicolelis. “Perceiving invisible light through a somatosensory cortical prosthesis”. In: Nature Communications 4 (2013), p. 1482. eprint: NIHMS150003.

[27] Eric E. Thomson et al. “Cortical neuroprosthesis merges visible and invisible light without impairing native sensory function”. In: eNeuro 4.6 (2017), pp. 1–17.

[28] Joseph E. O’Doherty, Solaiman Shokur, Leonel E. Medina, Mikhail A. Lebedev, and Miguel A.L. Nicolelis. “Creating a neuroprosthesis for active tactile exploration of textures”. In: Proceedings of the National Academy of Sciences of the United States of America 116.43 (2019), pp. 21821–21827.

[29] Maria C Dadarlat. “Artificial Sensory Feedback for Neural Prostheses”. PhD thesis. University of California, San Francisco, 2014, pp. 1078–1085.

[30] Maria C. Dadarlat and Philip N. Sabes. “Encoding and Decoding of Multi-Channel ICMS in Macaque Somatosensory Cortex”. In: IEEE Transactions on Haptics 9.4 (2016), pp. 508–514.

[31] Kevin A. Mazurek and Marc H. Schieber. “Injecting Instructions into Premotor Cortex”. In: Neuron 96.6 (2017), 1282–1289.e4.

[32] N A Fitzsimmons, W Drake, T L Hanson, M A Lebedev, and M A L Nicolelis. “Primate Reaching Cued by Multichannel Spatiotemporal Cortical Microstimulation”. In: Journal of Neuroscience 27.21 (2007), p. 5593.

[33] Amy L. Orsborn et al. “Closed-loop decoder adaptation shapes neural plasticity for skillful neuroprosthetic control”. In: Neuron 82.6 (2014), pp. 1380–1393.

[34] Richard A. Andersen, Tyson Aflalo, and Spencer Kellis. “From thought to action: The brain-machine interface in posterior parietal cortex”. In: Proceedings of the National Academy of Sciences of the United States of America 116.52 (2019), pp. 26274–26279.

[35] Sliman J. Bensmaia and Lee E. Miller. “Restoring sensorimotor function through intracortical interfaces: Progress and looming challenges”. In: Nature Reviews Neuroscience 15.5 (2014), pp. 313–325.

[36] Chethan Pandarinath and Sliman J Bensmaia. “The science and engineering behind sensitized brain-controlled bionic hands”. In: Physiological Reviews (2021).

[37] Maria C Dadarlat, Ryan A Canfield, and Amy L Orsborn. “Neural Plasticity in Sensorimotor Brain-Machine Interfaces”. In: Annual Review of Biomedical Engineering 15.2 (2023), pp. 51–76.

[38] Barry E Stein and Terrence R Stanford. “Multisensory integration: current issues from the perspective of the single neuron.” In: Nature reviews. Neuroscience 9.4 (2008), pp. 255–266.

[39] Sergejus Butovas and Cornelius Schwarz. “Spatiotemporal effects of microstimulation in rat neocortex: a parametric study using multielectrode recordings.” In: Journal of neurophysiology 90.5 (Nov. 2003), pp. 3024–39.

[40] Mark H Histed, Vincent Bonin, and R Clay Reid. “Direct activation of sparse, distributed populations of cortical neurons by electrical microstimulation.” In: Neuron 63.4 (2009), pp. 508–22.

[41] Michelle Armenta Salas et al. “Proprioceptive and cutaneous sensations in humans elicited by intra-cortical microstimulation”. In: eLife 7 (2018), pp. 1–11.

[42] Gregg a Tabot et al. “Restoring the sense of touch with a prosthetic hand through a brain interface.” In: Proc. Natl. Acad. Sci. U. S. A. 110.45 (2013), pp. 18279–84.

[43] Thierri Callier, Nathan W Brantly, Attilio Caravelli, and Sliman J Bensmaia. “The frequency of cortical microstimulation shapes artificial touch”. In: Proc Natl Acad Sci USA 117.2 (2020), pp. 1191–1200.

[44] Christopher Hughes and Takashi Kozai. “Dynamic amplitude modulation of microstimulation evokes biomimetic onset and offset transients and reduces depression of evoked calcium responses in sensory cortices”. In: Brain Stimulation 16 (2023), pp. 939–965.

[45] Isabelle A. Rosenthal et al. “Visual context affects the perceived timing of tactile sensations elicited through intra-cortical microstimulation”. In: bioRxiv (2024). eprint: https://www.biorxiv.org/content/early/2024/05/14/2024.05.13.593529.full.pdf.

[46] John S Choi et al. “Eliciting naturalistic cortical responses with a sensory prosthesis via optimized microstimulation”. In: Journal of Neural Engineering 13.5 (2016), p. 056007.

[47] Karthik Kumaravelu et al. “A comprehensive model-based framework for optimal design of biomimetic patterns of electrical stimulation for prosthetic sensation”. In: Journal of Neural Engineering 17.4 (2020), p. 046045.

[48] Katherine W. Scangos, Ghassan S. Makhoul, Leo P. Sugrue, Edward F. Chang, and Andrew D. Krystal. “State-dependent responses to intracranial brain stimulation in a patient with depression”. In: Nature Medicine 27.2 (2021), pp. 229–231.

[49] Maria C. Dadarlat, Yujiao Jennifer Sun, and Michael P. Stryker. “Activity-dependent recruitment of inhibition and excitation in the awake mammalian cortex during electrical stimulation”. In: Neuron 112.5 (2024), 821–834.e4.

[50] Richy Yun, Jonathan H. Mishler, Steve I. Perlmutter, Rajesh P. N. Rao, and Eberhard E. Fetz. “Responses of cortical neurons to intracortical microstimulation in awake primates”. In: eNeuro 10.4 (2023), ENEURO.0336–22.2023.

[51] John C. Tuthill and Eiman Azim. “Proprioception”. In: Current Biology 28.5 (2018), R194–R203.

[52] Ignacio Alonso et al. “Peripersonal encoding of forelimb proprioception in the mouse somatosensory cortex”. In: Nature Communications 14 (2023), p. 1866.

[53] Raeed H. Chowdhury, Joshua I. Glaser, and Lee E. Miller. “Area 2 of primary somatosensory cortex encodes kinematics of the whole arm”. In: eLife 9 (2020), pp. 1–31.

[54] Sanjiv K. Talwar et al. “Rat navigation guided by remote control”. In: Nature 417.6884 (2002), pp. 37–38.

[55] Andrew G. Richardson et al. “Learning active sensing strategies using a sensory brain–machine interface”. In: Proceedings of the National Academy of Sciences 116.35 (2019), p. 201909953.

[56] Konstantin Hartmann et al. “Embedding a panoramic representation of infrared light in the adult rat somatosensory cortex through a sensory neuroprosthesis”. In: Journal of Neuroscience 36.8 (2016), pp. 2406–2424.

[57] Hiroaki Norimoto and Yuji Ikegaya. “Visual cortical prosthesis with a geomagnetic compass restores spatial navigation in blind rats”. In: Current Biology 25.8 (2015), pp. 1091–1095.

[58] Maria C. Dadarlat, Joseph E. O’Doherty, and Philip N. Sabes. “A learning-based approach to artificial sensory feedback leads to optimal integration”. In: Nature Neuroscience 18.1 (2015), pp. 138–144.

[59] M J Prud’homme and J F Kalaska. “Proprioceptive activity in primate primary somatosensory cortex during active arm reaching movements.” In: Journal of neurophysiology 72.5 (1994), pp. 2280–301.

[60] Seungbin Park, Megan H. Lipton, and Maria C. Dadarlat. “Population-level encoding of somatosensation in mouse sensorimotor cortex”. In: bioRxiv (2025). eprint: https://www.biorxiv.org/content/early/2025/01/21/2025.01.21.634118.full.pdf.

[61] Stephanie Niklaus, Silvio Albertini, Tobias K. Schnitzer, and Nora Denk. “Challenging a myth and misconception: Red-light vision in rats”. In: Animals 10.3 (2020), pp. 1–14.

[62] Gary A Kane, Jonny L Saunders, and Alexander Mathis. “Real-time, low-latency closed-loop feedback using markerless posture tracking”. In: eLife 9 (2021), e61909.

[63] Nicholas J. Michelson, James R. Eles, Alberto L. Vazquez, Kip A. Ludwig, and Takashi D.Y. Kozai. “Calcium activation of cortical neurons by continuous electrical stimulation: Frequency dependence, temporal fidelity, and activation density”. In: Journal of Neuroscience Research 97.5 (2019), pp. 620–638.

[64] Maria C. Dadarlat, Yujiao Sun, and Michael P. Stryker. “Widespread activation of awake mouse cortex by electrical stimulation”. In: International IEEE/EMBS Conference on Neural Engineering, NER. Vol. 2019-March. 2019, pp. 1113–1117.

[65] Kevin C Stieger, James R Eles, Kip A Ludwig, and Takashi D Y Kozai. “Intracortical microstimulation pulse waveform and frequency recruits distinct spatiotemporal patterns of cortical neuron and neuropil activation”. In: Journal of Neural Engineering 19.2 (2022), p. 026024.

[66] Christopher P. Burgess et al. “High-Yield Methods for Accurate Two-Alternative Visual Psychophysics in Head-Fixed Mice”. In: Cell Reports 20.10 (2017), pp. 2513–2524.

[67] J. Sombeck et al. “Characterizing the short-latency evoked response to intracortical microstimulation across a multi-electrode array”. In: Journal of Neural Engineering 19 (2022), p. 026044.

[68] Jason M. Godlove, Erin O. Whaite, and Aaron P. Batista. “Comparing temporal aspects of visual, tactile, and microstimulation feedback for motor control”. In: Journal of Neural Engineering 11.4 (2014).

[69] Brandon M Ruszala and Marc H Schieber. “Injecting information in the cortical reach-to-grasp network is effective in ventral but not dorsal nodes”. In: Cell Reports 44 (2025), p. 115664.

[70] K. Novak, L. Miller, and J. Houk. “The use of overlapping submovements in the control of rapid hand movements”. In: Experimental Brain Research 144.3 (2002), pp. 351–364.

[71] Joseph G. Makin, Matthew R. Fellows, and Philip N. Sabes. “Learning Multisensory Integration and Coordinate Transformation via Density Estimation”. In: PLoS Computational Biology 9.4 (Apr. 2013), e1003035.

[72] Marc O. Ernst and Heinrich H. Bülthoff. “Merging the senses into a robust percept”. In: Trends in Cognitive Sciences 8.4 (2004), pp. 162–169.

[73] Jean Paul Noel and Dora E. Angelaki. “Cognitive, Systems, and Computational Neurosciences of the Self in Motion”. In: Annual Review of Psychology 73 (2022), pp. 103–129.

[74] Guido T. Meijer, Paul E.C. Mertens, Cyriel M.A. Pennartz, Umberto Olcese, and Carien S. Lansink. “The circuit architecture of cortical multisensory processing: Distinct functions jointly operating within a common anatomical network”. In: Progress in Neurobiology 174.December 2018 (2019), pp. 1–15.

[75] Jeffrey M. Yau, Gregory C. DeAngelis, and Dora E. Angelaki. “Dissecting neural circuits for multi-sensory integration and crossmodal processing”. In: Philosophical Transactions of the Royal Society B: Biological Sciences 370.1677 (2015), p. 20140203.

[76] Philip Coen, Timothy P.H. Sit, Miles J. Wells, Matteo Carandini, and Kenneth D. Harris. “Mouse frontal cortex mediates additive multisensory decisions”. In: Neuron 111 (2023), pp. 1–16.

[77] Michael L. Morgan, Gregory C. DeAngelis, and Dora E. Angelaki. “Multisensory Integration in Macaque Visual Cortex Depends on Cue Reliability”. In: Neuron 59.4 (Aug. 2008), pp. 662–673. eprint: NIHMS150003.

[78] R J van Beers, A C Sittig, and J J Gon. “Integration of proprioceptive and visual position-information: An experimentally supported model.” In: Journal of neurophysiology 81.3 (1999), pp. 1355–64.

[79] Kathleen E. Cullen. “The vestibular system: Multimodal integration and encoding of self-motion for motor control”. In: Trends in Neurosciences 35.3 (2012), pp. 185–196.

[80] Liping Yu, Benjamin a Rowland, and Barry E Stein. “Initiating the development of multisensory integration by manipulating sensory experience.” In: The Journal of Neuroscience 30.14 (Apr. 2010), pp. 4904–4913.

[81] Barry E Stein, Terrence R Stanford, and Benjamin A Rowland. “Development of multisensory integration from the perspective of the individual neuron”. In: Nature Reviews Neuroscience 15.8 (2014), pp. 520–535.

[82] M T Wallace, B N Carriere, T J Perrault, J W Vaughan, and B E Stein. “The development of cortical multisensory integration.” In: The Journal of neuroscience 26.46 (2006), pp. 11844–11849.

[83] J. Xu, L. Yu, B. a. Rowland, T. R. Stanford, and B. E. Stein. “Incorporating Cross-Modal Statistics in the Development and Maintenance of Multisensory Integration”. In: Journal of Neuroscience 32.7 (Feb. 2012), pp. 2287–2298.

[84] Edwin M. Maynard, Craig T. Nordhausen, and Richard A. Normann. “The Utah Intracortical Electrode Array: A recording structure for potential brain-computer interfaces”. In: Electroencephalography and Clinical Neurophysiology 102.3 (1997), pp. 228–239.

[85] Patrick T. Sadtler et al. “Neural constraints on learning”. In: Nature 512.7515 (2014), pp. 423–426.

[86] Matthew D Golub et al. “Learning by neural reassociation”. In: Nature Neuroscience 21 (2018), pp. 607–616.

[87] Emily R Oby et al. “New neural activity patterns emerge with long-term learning”. In: Proceedings of the National Academy of Sciences of the United States of America 116.30 (2019), pp. 15210–15215.

[88] Karthik Kumaravelu and Warren M. Grill. “Neural mechanisms of the temporal response of cortical neurons to intracortical microstimulation”. In: Brain Stimulation 17.2 (2024), pp. 365–381.

[89] Taylor G. Hobbs et al. “Biomimetic stimulation patterns drive natural artificial touch percepts using intracortical microstimulation in humans”. In: Journal of Neural Engineering 22.3 (2025).

[90] Charles M. Greenspon, Natalya D. Shelchkova, Taylor G. Hobbs, Sliman J. Bensmaia, and Robert A. Gaunt. “Intracortical microstimulation of human somatosensory cortex induces natural perceptual biases”. In: Brain Stimulation 17.6 (2024), pp. 1178–1185.

[91] Jacob L. Segil, Ivana Cuberovic, Emily L. Graczyk, Richard F.ff Weir, and Dustin Tyler. “Combination of Simultaneous Artificial Sensory Percepts to Identify Prosthetic Hand Postures: A Case Study”. In: Scientific Reports 10.1 (2020), pp. 1–15.

[92] Charles M. Greenspon et al. “Evoking stable and precise tactile sensations via multi-electrode intra-cortical microstimulation of the somatosensory cortex”. In: Nature Biomedical Engineering 9.6 (2025), pp. 935–951.

[93] Yu Cheng Pei, Steven S. Hsiao, James C. Craig, and Sliman J. Bensmaia. “Shape invariant coding of motion direction in somatosensory cortex”. In: PLoS Biology 8.2 (2010), e1000305.

[94] T. D. Albright. “Direction and orientation selectivity of neurons in visual area MT of the macaque”. In: Journal of Neurophysiology 52.6 (1984), pp. 1106–1130.

[95] Jacob Reimer et al. “Pupil Fluctuations Track Fast Switching of Cortical States during Quiet Wakefulness”. In: Neuron 84.2 (2014), pp. 355–362.

[96] Joseph T. Sombeck and Lee E. Miller. “Short reaction times in response to multi-electrode intracortical microstimulation may provide a basis for rapid movement-related feedback”. In: Journal of Neural Engineering 17 (2020), p. 016013.

[97] Anne E. Urai et al. “Citric acid water as an alternative to water restriction for high-yield mouse behavior”. In: eNeuro 8.1 (2021), pp. 1–8.

