## Supplementary material for "Rapid integration of artificial sensation": Suppl_Figures_1_and_2

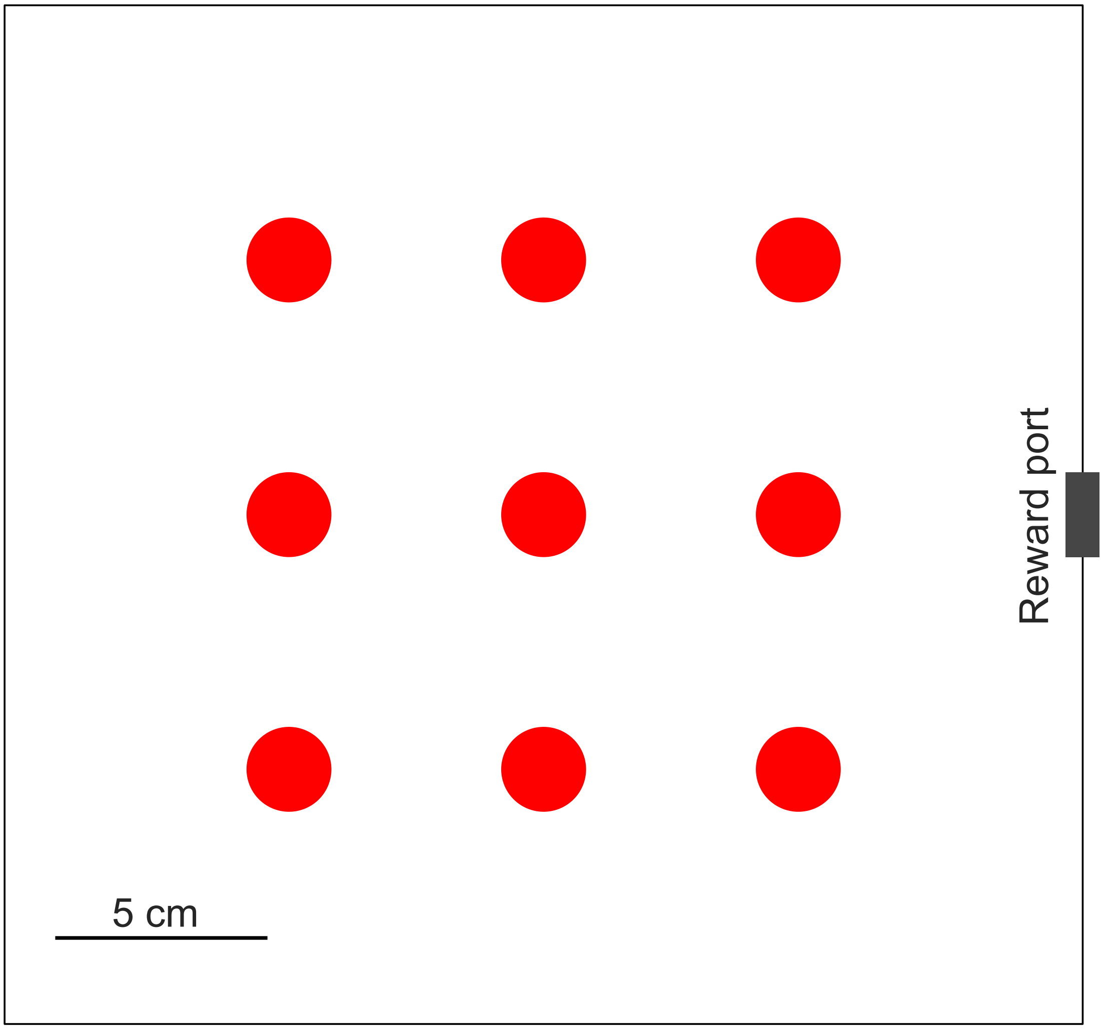


Figure S 1: Behavioral targets. **A.** Schematic showing the nine target locations upon the cage floor (aerial view). Each target is 2 cm wide, created by illuminating units on an LED matrix placed below the clear cage floor. A reward port sits on the center of the far right wall. Targets are spaced 6 cm apart and are chosen randomly per trial.

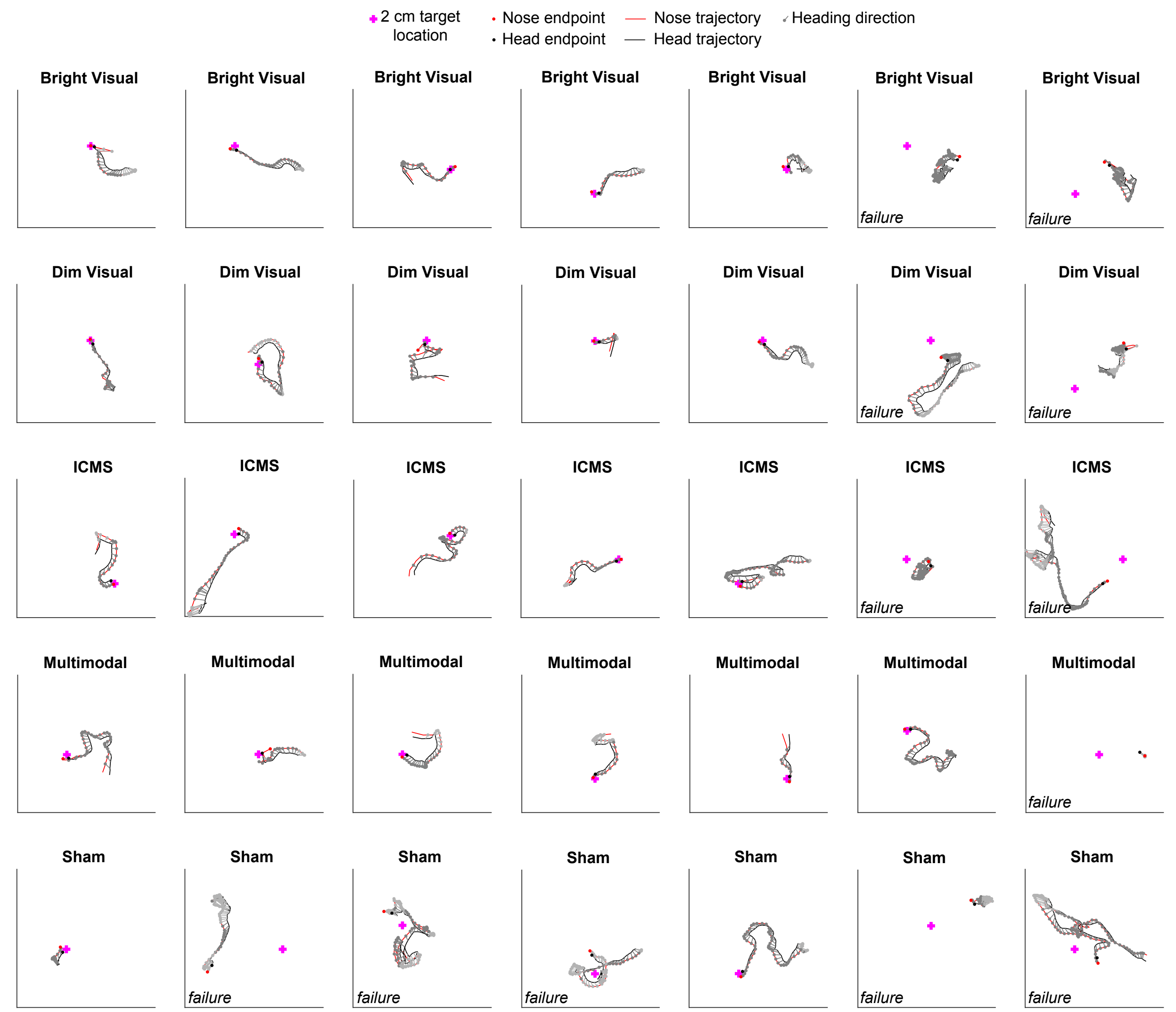


Figure S 2: Sample single-trial trajectories from a single mouse during the last five behavioral sessions. In addition to showing the mouse's nose and head position during the trial (red and black traces), the calculated heading is shown every 10 samples, where the circle denotes nose position. The heading is colored to show when the mouse is "facing" the target (dark gray) vs. not facing the target (light gray), where facing is defined as having the target location within eyesight — a field of view of $246.8^o$ centered on the mouse's heading Samonds et al., 2019. In the analysis in the main text, the trials are cropped to the exclude any movements before the mouse was facing the target. The target (invisible on ICMS and sham trials) is shown here as a magenta cross. The target's true size is 2 cm in diameter, shaped as a circle constructed from an illuminated LED matrix; here is it shown to scale relative to the sides of the cage (black lines).
